# Critical Limb Ischemia Induces Remodeling of Skeletal Muscle Motor Unit and Myonuclear- and Mitochondrial-Domains

**DOI:** 10.1101/426403

**Authors:** Mahir Mohiuddin, Nan Hee Lee, June Young Moon, Woojin M. Han, Shannon E. Anderson, Jeongmoon J. Choi, Shadi Nakhai, Thu Tran, Berna Aliya, Do Young Kim, Aimee Gerold, Laura Hansen, W. Robert Taylor, Young C. Jang

## Abstract

Critical limb ischemia, the most severe form of peripheral artery disease, leads to extensive damage and alterations in skeletal muscle homeostasis. Although recent developments towards revascularization therapies have been introduced, there has been limited research into treatments for ischemic myopathy. To elucidate the regenerative mechanism of the muscle stem cell and its niche components in response to ischemic insults, we explored interactions between the vasculature, motor neuron, muscle fiber, and the muscle stem cell. We first investigated changes in the neuromuscular junction and motor neuron innervation following a surgical hindlimb ischemia model of critical limb ischemia in mice. Along with previous findings that support remodeling of the neuromuscular junction, we report that ischemic injury also causes significant alterations to the myofiber through a muscle stem cell-mediated increase of myonuclei number per fiber, a concomitant decrease in myonuclear domain size, and an increase in relative mitochondrial content per myonucleus. These results indicate that as a regenerative response to critical limb ischemia, myofibers exhibit myonuclear expansion to allow enhanced transcriptional support and an increase in mitochondrial content for bioenergetic need of the energy-demanding tissue regeneration.

## 1. Introduction

Peripheral artery disease (PAD) is a degenerative cardiovascular disease characterized by abnormal perfusion in the limbs due to occlusions of the blood vessels. In its most severe form, referred to as critical limb ischemia (CLI), where blood flow is insufficient to maintain tissue viability, amputation rates can reach up to 50% [1]. While recent regenerative medicine approaches on collateral vessel formation have made some progress, the myopathy and dysregulation of the skeletal muscle in CLI have not been thoroughly investigated [2-4]. Recent evidence suggests that the ischemia-induced loss of viable muscle tissue, which acts as an indispensable matrix of growth factors and biomechanical support for new vessel formation, may be the cause of failure for current revascularization therapies [5, 6]. This paradox emphasizes the importance of skeletal muscle tissue as a therapeutic target in conjunction with angiogenic therapies in PAD.

Skeletal muscles possess a remarkable regenerative capacity due to the presence of endogenous resident muscle stem cells (MuSCs), also known as satellite cells [7]. Upon injury, Pax7^+^ quiescent MuSCs, which reside adjacent to basal lamina and sarcolemma of the multinucleated myofiber, undergo asymmetric division in which committed progenies proliferate, differentiate, and fuse with existing myofibers or form *de novo* myofibers, while other populations of MuSC progeny self-renew to replenish the quiescent stem cell pool for future rounds of regeneration [8]. Emerging evidence suggests that MuSC function and regenerative capacity are dictated by cellular and acellular interactions with its surrounding microenvironment, or niche, such as the vasculature, neuromuscular junction (NMJ) of a motor neuron, myofibers, interstitial cells of the extracellular matrix, as well as various infiltrating immune cells [9]. Of these niche components, the vasculature and microvessels are located close to MuSCs and provide necessary oxygen, nutrients, and growth factors required for MuSC function and muscle homeostasis. In addition, vascular networks near MuSCs play a crucial role in recruiting circulating stem cells and transporting immune cells during the initial phase of muscle repair [10, 11]. Thus, the lack of functional blood perfusion to skeletal muscle not only disrupts muscle homeostasis by limiting oxygen and nutrient delivery but also compromises muscle regeneration [12].

It has been well documented that motor neuron innervation maintains muscle homeostasis by regulating excitation-contraction coupling as well as controlling the gene expression pattern of myofibers [13]. Denervation of the muscle fiber due to injury or neuromuscular disease results in muscle wasting and remodeling of motor units [14, 15]. Anatomically, peripheral nerves, such as lower motor neurons, are located close in contact with vasculature (neurovascular congruency) [16] and exhibit functional interdependency. In support of this notion, some of the axonal guidance factors are known to possess angiogenic properties [17], and other well-known angiogenic factors, such as vascular endothelial growth factor (VEGF), guide Schwann cell-mediated peripheral nerve regeneration [18]. As such, when blood flow is restricted in ischemic injury, motor neurons undergo rapid Wallerian degeneration, and their regenerative response is activated [19, 20]. Moreover, a recent report showed that the regeneration of the motor neuron and its corresponding neuromuscular synapses are in part linked to an increase in activation of MuSC and myogenesis [21]. Conversely, a genetic depletion of MuSCs diminishes the regenerative response of the neuromuscular junction [22], demonstrating crosstalk between the NMJ and MuSC. Collectively, the majority of previous studies reported tissue-specific interaction in response to ischemic insults (i.e., ischemia – muscle stem cell, ischemia – muscle fiber, ischemia – motor neuron) and an investigation elucidating the mechanistic crosstalk between collateral vascularization and angiogenesis, motor neuron regeneration, and muscle stem cells and muscle regeneration altogether has not been conducted. Therefore, in this study, we aim to characterize the temporal interactions between muscle stem cells, myofibers, and motor units in response to ischemia/reperfusion injury. To this end, we explored remodeling of the MuSC niche components, notably the motor neuron, NMJ, and myofiber, at various time points to elucidate the sequential regeneration of the tissues following limb claudication. Using the murine hindlimb ischemia (HLI) model that manifests similar pathophysiology as human CLI, we demonstrated ischemia-induced early necrotic deterioration of muscle fibers and motor neuron/NMJ, followed by regeneration of both tissues that persists for at least 56 days, and surprising alterations to the myonuclear and mitochondrial networks within the myofiber. The myonuclear domain, defined as the cytoplasmic volume of myofiber transcriptionally governed by each nucleus, was quantified to characterize the myonuclear network. In effect, this paper highlights the highly orchestrated remodeling of the MuSC and its niche components following CLI and provides a basis to investigate multi-scale therapies.

## 2. Materials and Methods

### 2.1. Animal models

All animal procedures were conducted under the approval of the Institutional Animal Care and Use Committee (IACUC) of Georgia Institute of Technology. All mice in this study were either *C57BL/6J* genetic background or backcrossed with *C57BL/6J* for more than 6 generations and were initially purchased from Jackson Laboratory. Mice were bred and maintained in pathogen free conditions with 12-12 light/dark cycle in the Physiological Research Laboratory (PRL) at Georgia Institute of Technology. For muscle stem/satellite cell reporter, mice expressing a tamoxifen-inducible Cre from the endogenous Pax7 promoter were bred with mice carrying a loxP-flanked STOP cassette followed by tdTomato in the ROSA26 locus. Other transgenic reporter mice include: *Thy1*-*EYFP (line 16)* for motor neuron [23], *mitoDendra2* for mitochondria [24, 25], and *PV Cre^ER^; ChR2*-*EYFP* (tamoxifen-inducible Cre recombinase inserted into PV locus bred with mice carrying a loxP-flanked STOP cassette followed by ChR2-EYFP) for expression of EYFP driven by calcium binding protein parvalbumin to detect type IIB and IIX fast-twitch fibers [26]. Both male and females aged between 3 and 6 months were used in a randomized manner for all experiments in this study.

### 2.2. Surgical procedure

To study the effects of critical limb ischemia, we employed a well-characterized murine hindlimb ischemia (HLI) surgical ligation model [27]. Briefly, the femoral artery and vein were ligated with sutures between the superficial epigastric artery and profunda femoris artery. A second ligation was made proximal to the branching of the tibial arteries and the segment of vessels between the two ligations was excised. The sham surgery, where the femoral artery and vein were exposed similar to the method above without ligation or excision, was performed on the contralateral leg. Animals were maintained for 3-56 days following HLI and Laser Doppler Perfusion Imaging (LDPI) was performed on a MoorLDI Imager before euthanization by CO_2_ inhalation.

### 2.3. Histochemistry and immunostaining

Immediately following euthanization of animals, the tibialis anterior (TA) muscles were either snap frozen for cryo-sectioning or fixed in 4% paraformaldehyde for single myofiber isolation. Frozen TA was sliced into 10 μm sections while fixed TA was mechanically separated into 20-30 single myofibers from random areas of the muscle. The gastrocnemius was frozen in liquid nitrogen for Western blot analysis while the EDL was fixed for immunostaining. Hematoxylin and eosin (H&E) and immunofluorescence were performed as previously described [28]. Antibodies can be found in Supplemental Table 1 for dilution factors, vendors, and catalog numbers of materials. All images were taken on either Zeiss Axio Observer D1 or Zeiss 700 Laser Scanning Confocal microscopes and quantified using ImageJ.

### 2.4. Mitochondria Functional Testing

Mitochondria were isolated from lower limb muscles distal to the knee by differential centrifugation. Muscles were digested with dispase II and trypsin, homogenized, centrifuged at 12,000 *g* for 10 minutes, and resuspended in Chappel-Perry buffer II (see Supplementary Table 2 for detail). The suspensions were then centrifuged at 600 *g* for 10 minutes and the supernatants centrifuged at 7,000 *g* to pellet the mitochondria. 300 μg of mitochondria resuspended in respiration buffer were used immediately in the Oroboros Oxygraph-2k high resolution respirometer for basal (state 1) oxygen consumption rate and H_2_O_2_ generation detected with the Amplex Red assay kit.

### 2.5. Western Blot Analysis

Homogenized gastrocnemii were normalized to total protein concentration, run through 4–20% Criterion TGX Gels (Bio-Rad 5671093), and transferred using a Trans-Blot Turbo System. Antibodies used can be found in Supplementary Table 1. Membranes were imaged on Li-Cor Odyssey CLx-1050 Infrared Imaging System and bands were quantified on Li-Cor Image Studio V5.2.

### 2.6. Statistical Analysis

All statistical analyses in this study were performed on GraphPad Prism 7 and data is presented as mean ± standard deviation (SD). Data from the contralateral controls did not change over time, therefore multiple comparison tests for significance were performed between mean of control and each timepoint. For experiments in which data was collected from different mice over time, a one-way ANOVA with Tukey’s *post hoc* test was used. A paired two-tailed t-test was used to compare the injured hindlimbs to their contralateral controls. A *p*-value of less than 0.05 was considered statistically significant.

## 3. Results

### 3.1. Hindlimb ischemia induces skeletal muscle regeneration

To investigate morphological changes to the skeletal muscle fibers and muscle stem cell niche following chronic hindlimb ischemia (HLI), we first correlated blood perfusion to muscle regeneration. Laser Doppler perfusion imaging was used to quantify the abnormal perfusion in the ischemic leg (right-hand side of each scan) to demonstrate a significant decrease in perfusion to the ischemic leg compared to its contralateral control (left-hand side of each scan) at all timepoints up to 56 days following HLI (Fig. 1A and Supplementary Fig. S1A). As expected, hematoxylin-eosin (Fig. 1B) and immunofluorescent staining (Fig. 1C) of cross-sections from the tibialis anterior muscle (TA) showed extensive myofiber damage and presence of necrotic fibers, markedly at day 3 and 7. A high number of nuclei within the interfibrillar space was also observed at 7 days following HLI, implying immune cell infiltration required for clearance of dead or damaged myofibers [11]. Reperfusion to the ischemic limb did not improve until day 14 (Supplementary Fig. S1B), where we found a coinciding peak in embryonic myosin heavy chain (eMHC) positive myofibers (Fig. 1D), a marker for early stages of regeneration. This suggests that sufficient perfusion through collateral vascularization parallels activation of muscle stem cells and muscle regeneration. To further assess myofiber regeneration, we noted a substantial increase in percentage of centrally nucleated fibers, characteristic of regenerating myofibers, at day 14 that persisted for up to 56 days (Fig. 1F). Furthermore, even though muscle fibers are regenerating, muscle atrophy, as measured by overall fiber cross sectional area, did not fully recover until day 56, suggesting that the functional deficiency may be prolonged following ischemic myopathy (Fig. 1E). In accordance with other models of ischemia [12], we observed a delayed regenerative response to ischemia compared to chemical modes of injury such as cardiotoxin, notexin, barium chloride, and glycerol, where immune cell infiltration was seen at days 3-4 and both eMHC^+^ and centrally nucleated fibers were decreased by day 14 following injury [29, 30]. These data show a delayed ischemia-induced regenerative response of skeletal muscle tissue, dependent on reperfusion to the hindlimb, that continues for at least 56 days.

**Figure 1.**
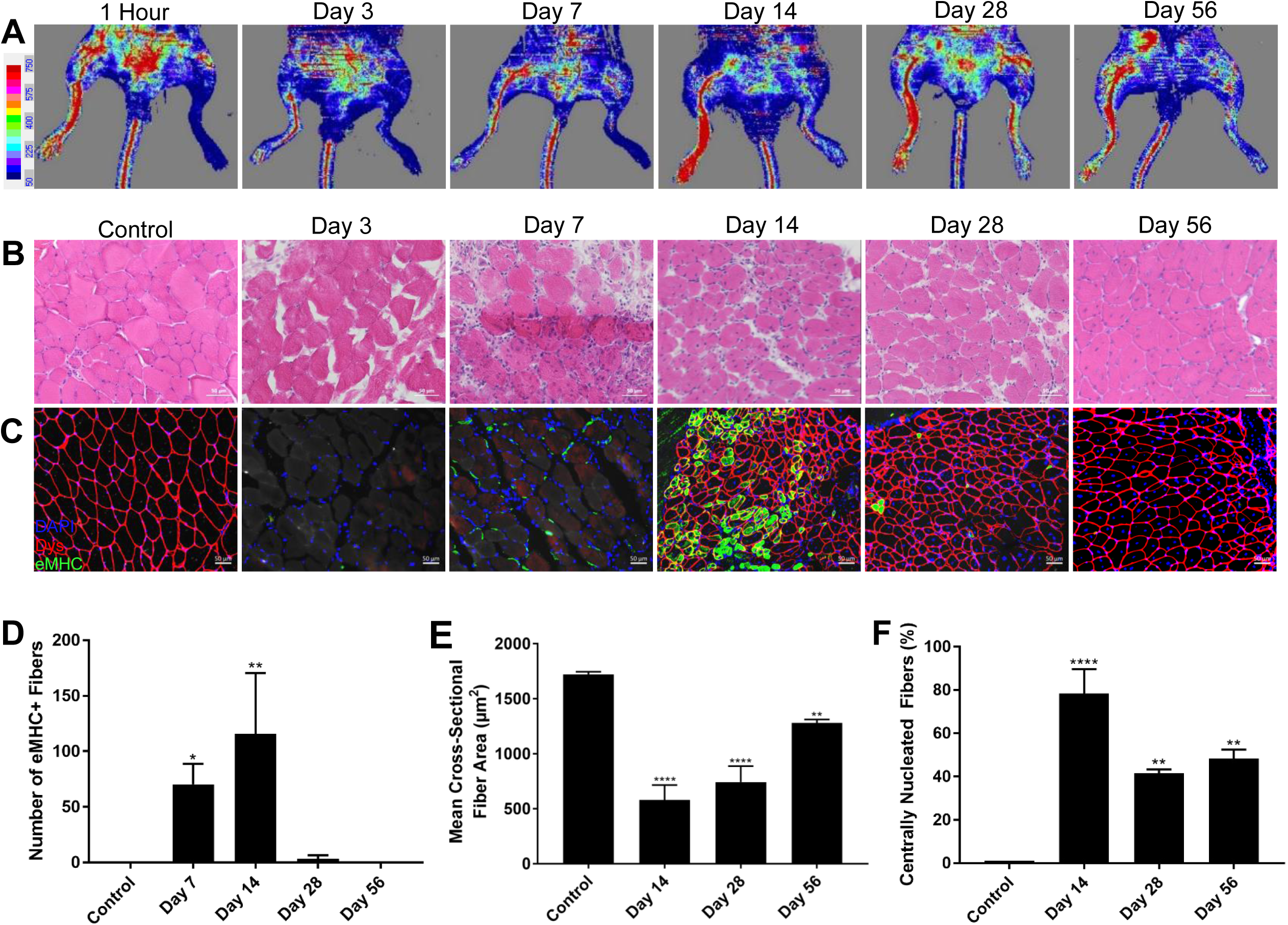
Characterization of CLI mouse model and skeletal muscle regeneration. (A) LDPI of ventral mouse hindlimbs 1 hour, 3 days, 7 days, 14 days, 28 days, and 56 days following HLI. Control leg on left, ischemic leg on right. Scale bar represents blood flow perfusion by color. (B) H&E staining of TA cross-sections in control, and 3 days, 7 days, 14 days, 28 days, and 56 days following HLI. (C) Immunohistochemistry of TA cross-sections in control, and 3 days, 7 days, 14 days, 28 days, and 56 days following HLI. Dystrophin pseudo-colored in red, eMHC in green, nuclei in blue. Scale bars on cross-sections represent 50 μm. (D) Total number of eMHC+ fibers within a 0.33 mm^2^ field of view for control, 7 days, 14 days, and 56 days following HLI. (E) Mean cross-sectional fiber area of 4 random 0.33 mm^2^ fields of view of the TA using dystrophin in control, 14 days, 28 days, and 56 days following HLI. (F) Percentage of centrally nucleated fibers in control, 14 days, 28 days, and 56 days following surgery. n=3, ^*^*p*<0.05, ^**^*p*<0.01, ^***^*p*<0.001, ^****^*p*<0.0001 compared to control.

### 3.2. Hindlimb ischemia induces skeletal muscle motor unit remodeling

In the next set of experiments, we examined the effects of ischemia on motor neuron disruption and subsequent denervation at the neuromuscular junction (NMJ). Given that a decline in innervation and loss of motor units are known causes of muscle degeneration [31], we tested whether denervation plays a role in the ischemia-induced atrophic process. Using Thy1-YFP reporter mice for YFP expression driven by the *Thy1* regulatory element [23], we observed marked alterations in NMJs in the ischemic extensor digitorum longus muscle (EDL) compared to control (Fig. 2A). Notably, at days 3 and day 7, presynaptic terminals of the motor axons exhibited abnormal thinning and characteristics of Wallerian degeneration while the postsynaptic endplates were fragmented compared to the normal pretzel-like morphology seen at day 0. When muscle fibers are innervated, the presynaptic nerve terminal and postsynaptic AChR of the NMJ overlay each other. Thus, to quantify innervation states of the muscle fibers, NMJs of each group were categorized as normal NMJ morphology (pretzel-like, convoluted folding structure with 100% overlay), partially denervated (fragmented AChR with between 25% and 99% overlay), or completely denervated (fragmented AChR with less than 25% overlay). Representative images of denervation states are shown in Supplemental Fig. S2A. Following HLI, we report a decrease in NMJs with normal morphology and an increase in both partial and complete denervation in ischemia-affected muscle compared to day 0 at all subsequent timepoints, with peak denervation observed at days 3 and 7. Interestingly, denervation of post-synaptic muscle lasted until at least day 56 (Fig. 2C). These data further suggest persistent functional deficiency in myofiber excitation-contraction coupling and corroborates the continued regeneration of skeletal muscle tissues up to 56 days.

**Figure 2.**
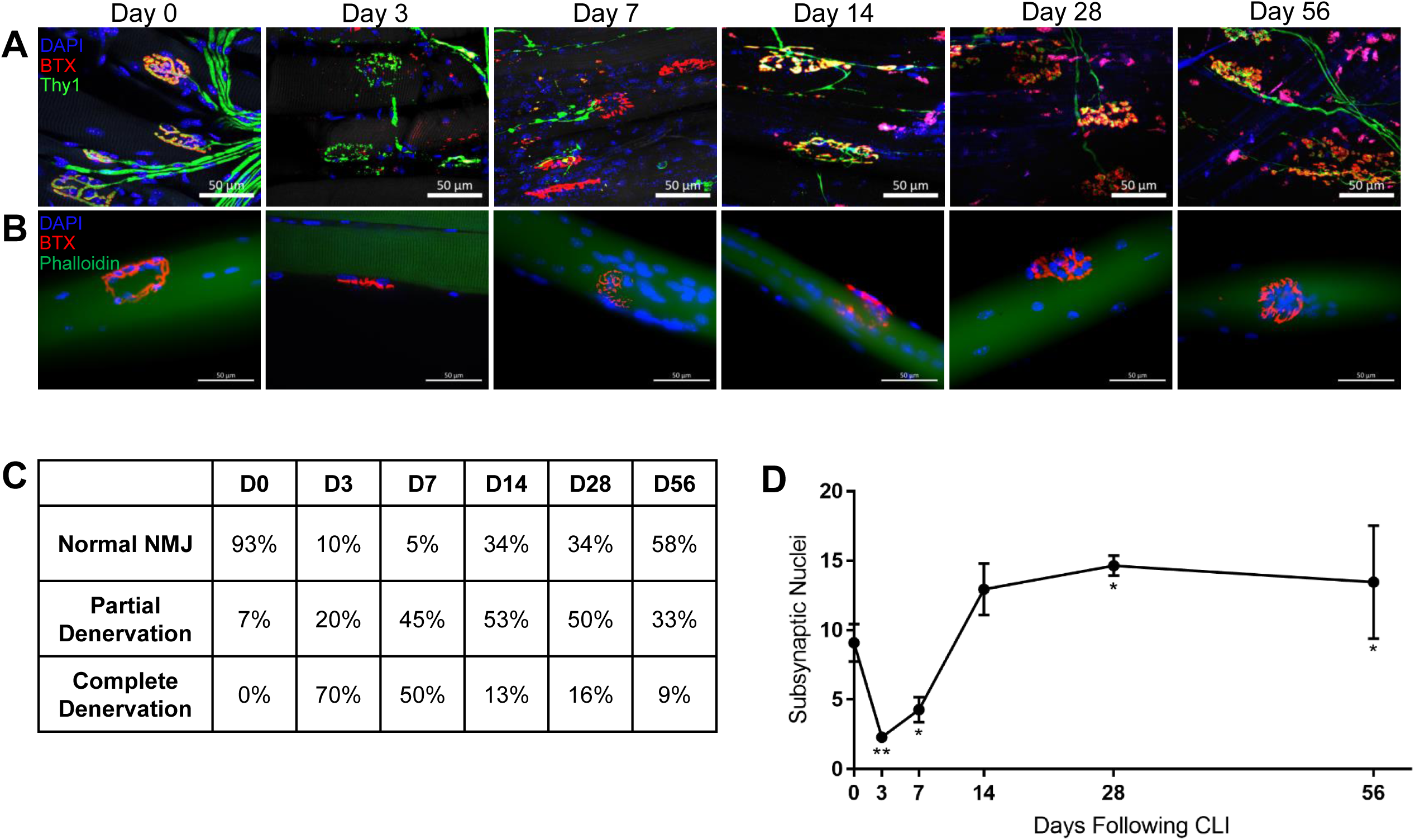
Remodeling of the motor unit. (A) NMJ in EDL of 1 hour (day 0), 3 days, 7 days, 14 days, 28 days, and 56 days following HLI. Nuclei pseudo-colored in blue, α-bungarotoxin in red, Thy1 in green. Maximum intensity projection was performed on images from confocal microscopy. (B) NMJ on single myofibers from TA from 1 hour (day 0), 3 days, 7 days, 14 days, 28 days, and 56 days following HLI. Nuclei pseudo-colored in blue, α-bungarotoxin in red, phalloidin in green. (C) Table of percentages of normal, partially denervated, and completely denervated NMJ in EDL of 1 hour (day 0), 3 days, 7 days, 14 days, 28 days, and 56 days following HLI. (D) Number of subsynaptic myonuclei within each NMJ of single myofibers from TA. All scale bars represent 50 μm. n=3, ^*^*p*<0.05, ^**^*p*<0.01 compared to day 0.

Next, we performed single myofiber analyses randomly isolated from TA muscle of control and HLI, comprised of almost all fast-twitch oxidative-glycolytic (type IIA) and fast-twitch glycolytic (types IIB/X) with few slow-twitch oxidative (type I) fibers [32], to reveal any remodeling of the NMJ endplates at the individual muscle fiber level. It is worth noting that since fiber type is in part determined by innervation from a specific motor neuron type, alteration in motor unit organization may lead to a shift in fiber type [33]. Indeed, we observed an ischemia-induced loss of slow-twitch type I fibers 14 and 28 days following injury while percentages of fast-twitch type IIA and types IIB/X fibers were unchanged (Supplementary Fig. S2B, C, D). Regardless, due to the fragmentation of acetylcholine receptors observed in isolated myofibers (Fig. 2B), we counted the subsynaptic nuclei that are responsible for maintenance of the NMJ through regulation of gene expression [34]. Despite a transient decrease in subsynaptic nuclei per NMJ at days 3 and 7, which may be attributed to fragmented NMJs in denervated myofibers [35], we observed an unexpected increase in the number of subsynaptic nuclei 28 and 56 days following HLI compared to day 0 (Fig. 2D). The increased subsynaptic nuclei coincide with the regeneration of the NMJ, indicating a dependence of NMJ restoration on the number of subsynaptic nuclei. Therefore, results suggest that ischemia-induced denervation plays a role in muscle atrophy and the remodeling of subsynaptic nuclei assist in the repair of denervated NMJ for up to 56 days following HLI.

### 3.3. Myonuclear domain decreases following hindlimb ischemia

To further elucidate the role of MuSCs and myogenesis in the regenerative process of the ischemic myopathy, we examined changes to myonuclear content induced by ischemia. To achieve this, single myofibers were isolated (Fig. 3A) and as seen in immunohistochemistry images, ischemic fibers were centrally nucleated. Myonuclei were assembled into longitudinal rows along the center of the myofiber, notably at days 7 and 14, although these rows migrate towards the periphery at 28 and 56 days following HLI. Surprisingly, despite the degeneration of muscle tissue following ischemic injury, a substantial increase in myonuclei number was observed at day 7 that prevails for at least 56 days (Fig. 3B). In addition, we assessed alterations in myonuclear domain, the cytoplasmic volume of myofiber governed by each myonucleus. Intriguingly, the size of the myonuclear domain was significantly decreased at 7 and 14 days following injury (Fig. 3C), likely as a regenerative response to ischemia in order to enhance transcriptional regulation of the myofiber cytoplasm within its domain, but returns to normal after the tissue perfusion was restored. Since myofibers are multinucleated syncytia formed by fusion of the differentiated myotubes [8], we then investigated the frequency of MuSCs following ischemia to determine the source of the accreted myonuclei in the MuSC niche. To explore MuSC frequency, we isolated single myofibers from a tamoxifen-inducible MuSC reporter mouse we generated by crossing *Pax7 Cre^ER^* with *ROSA26*-*tdTomato* mice (Fig. 3D). As hypothesized, we report a substantial increase in Pax7^+^ MuSC content at days 7 and 14 following ischemia, with a peak at day 7 that accompanies the accretion of myonuclei and concomitant decrease in myonuclear domain (Fig. 3E). Because Pax7 is a canonical marker for quiescent MuSC [8], the greater frequency of Pax7^+^ cells imply that MuSCs have undergone adult myogenesis and asymmetric division to increase self-renewal of MuSCs following ischemia to replenish the pool of quiescent MuSCs to continue regenerating and for future rounds of regeneration. Furthermore, the increased number of myonuclei also indicates active differentiation and fusion events of MuSCs into the myofiber following claudication of muscle. Taken together (Fig. 3F), these data demonstrate that HLI muscle exhibit a considerable remodeling of myonuclear domain possibly to enhanced transcriptional control of the newly formed myofiber.

**Figure 3.**
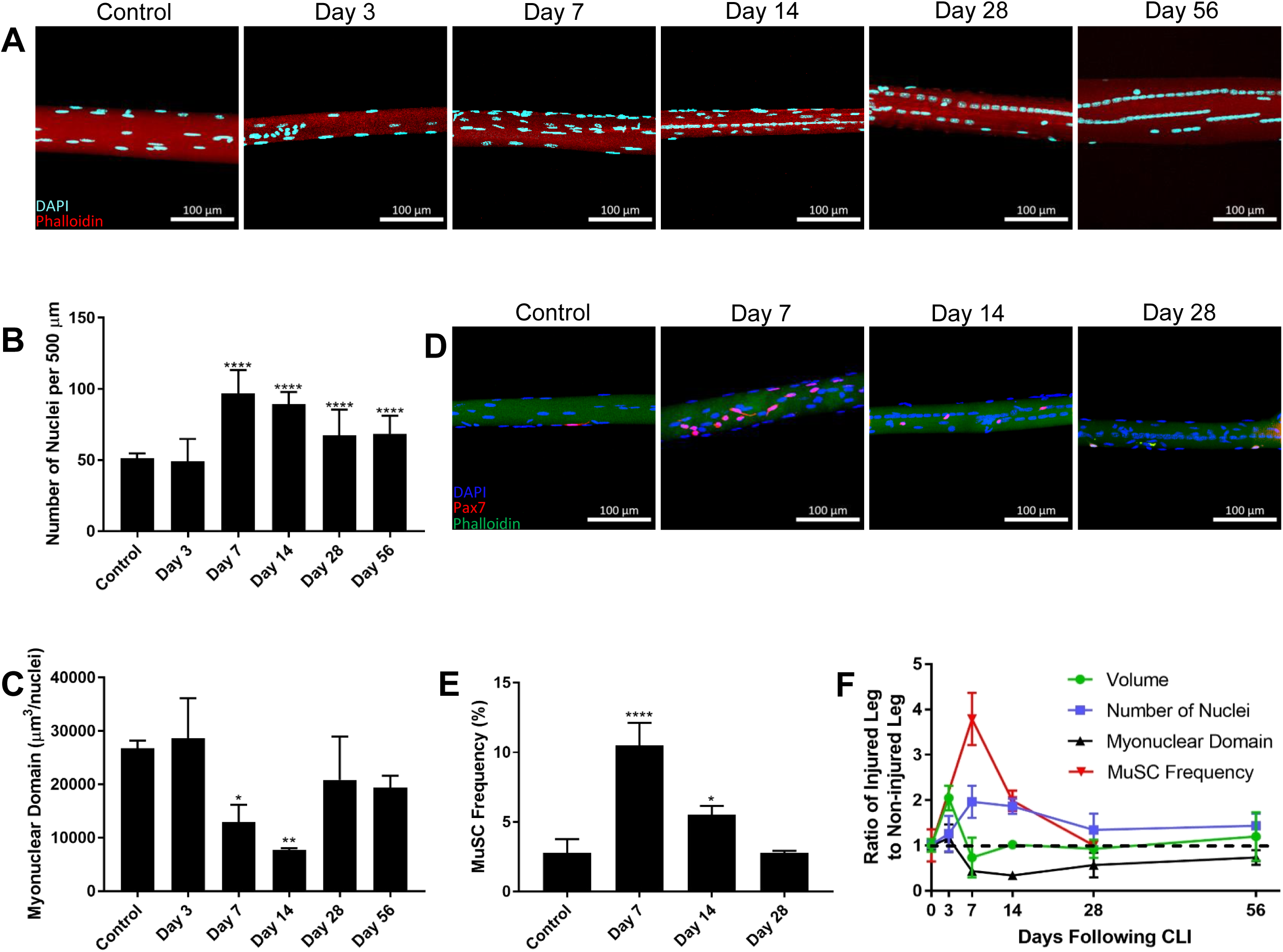
Changes in myonuclear domain following CLI. (A) Z-stack confocal imaging of single myofibers from TA of control, 3 days, 7 days, 14 days, 28 days, and 56 days following HLI. Nuclei pseudo-colored in light blue, phalloidin in red. (B) Z-stack confocal imaging of single myofibers from TA of Pax7-TdTomato mice control, 7 days, 14 days, and 28 days following HLI. Nuclei pseudo-colored in blue, Pax7 in red, phalloidin in green. All scale bars represent 100 μm. Maximum intensity projection performed on all z-stack images. (C) Number of myonuclei per 500 μm of myofiber at various timepoints. (D) Myonuclear domain of single myofibers at various timepoints calculated as myofiber volume divided by number of myonuclei. (E) MuSC frequency of single myofibers at various timepoints reported as percentage of Pax7^+^ nuclei out of total nuclei. n=3, ^*^*p*<0.05, ^**^*p*<0.01, ^****^*p*<0.0001 compared to control. (F) Myofiber volume, myonuclei number, myonuclear domain, and MuSC frequency as a ratio of ischemic to control myofiber. Dashed line represents a ratio of 1.

### 3.4. Mitochondrial domain increases following hindlimb ischemia

Another critical component of muscle repair is the bioenergetics of regenerating myofibers. Muscle regeneration is a highly energy dependent process and an increase in mitochondrial volume usually couples with adult myogenesis. Moreover, since mitochondrial DNA accounts for only 13 of the proteins that constitute mitochondria while over 99% of mitochondrial genes are nuclear-encoded [36], we examined changes in mitochondrial to myonuclear ratio. To this end, we isolated single myofibers from reporter mice expressing a mitochondrial-targeted fluorescent protein (Dendra2) [25] and imaged the myofibers on a confocal microscope with constant gain and laser power to delineate relative mitochondrial volume of the myofiber (Fig. 4A). In control samples, we noted local mitochondrial networks as punctate and compartmentalized into columns along the z-line of the myofiber contractile apparatus. At 7 days post-HLI, however, mitochondrial networks in ischemic myofibers spanned several sarcomeres (Fig. 4A) [24]. The reticulum of mitochondria are observed again at 14 and 28 days following HLI, although high mitochondrial densities with abnormal organization are also found between the central nuclei, indicated by the red arrowheads (Fig. 4A). We then quantified mitochondrial domain, or relative mitochondrial volume per nucleus, as a metric to describe changes in the number of mitochondria as an effect of the myonuclear domain size. Surprisingly, no change in mitochondrial domain is detected at 7 and 14 days following ischemia when myonuclear domain is at its smallest. Rather, there is a subsequent increase in mitochondrial domain 28 days following HLI, after myonuclear domain size has returned to normal but myonuclei number is still increased (Fig. 4B). The increased mitochondrial content at 28 days insinuates greater mitochondrial biogenesis in regenerating myofibers demarcated near the centrally located myonuclei.

**Figure 4.**
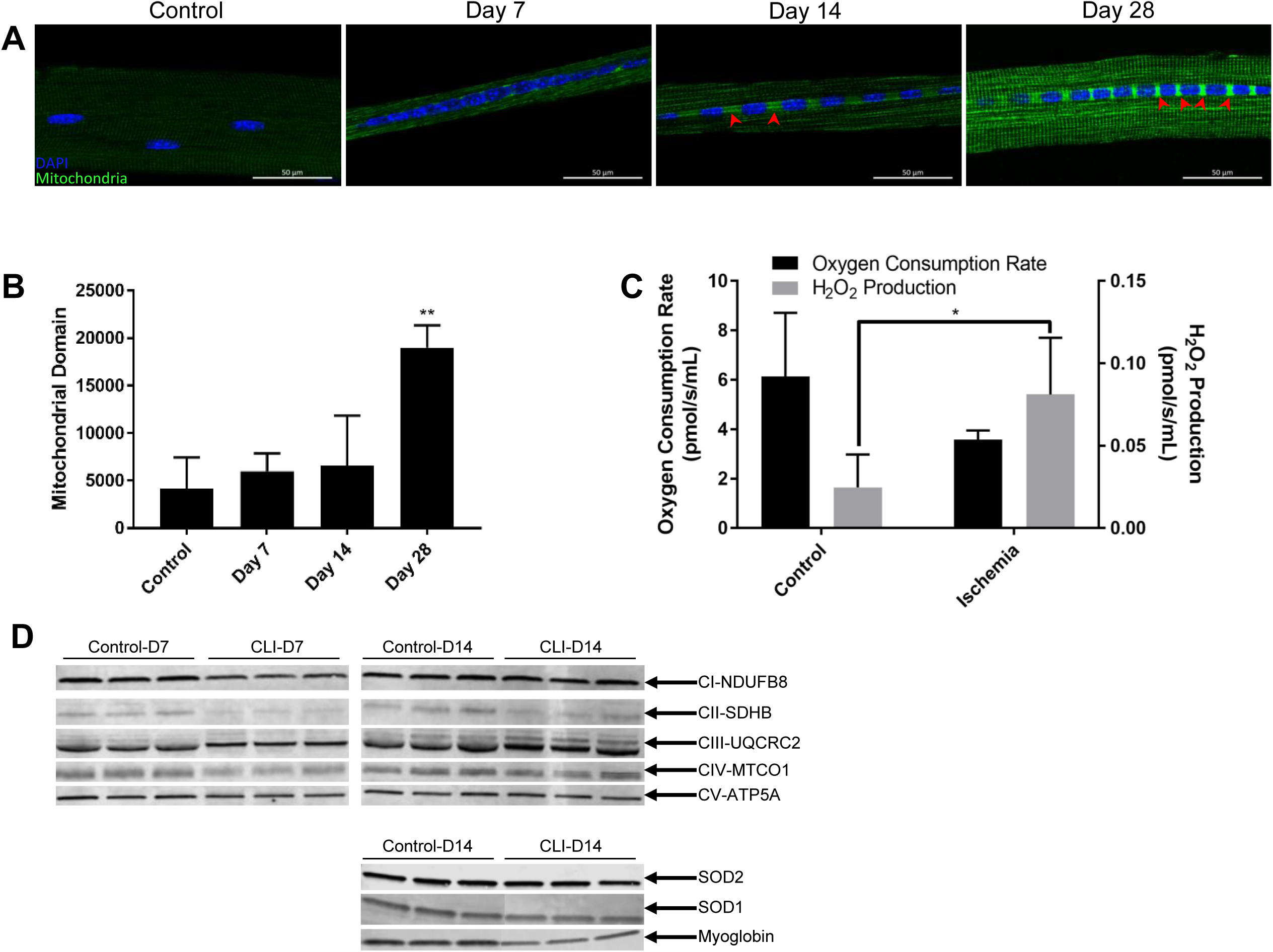
Changes in mitochondrial domain following CLI. (A) Confocal imaging of single myofibers from TA of mitoDendra2 mice control, 7 days, 14 days, and 28 days following HLI using constant gain and laser power. Nuclei pseudo-colored in blue, mitochondria in green. Red arrowheads indicate areas of high mitochondrial density. Scale bar represents 20 μm. (B) Mitochondrial domain of single myofibers at various timepoints calculated as relative integrated fluorescent density of mitochondria divided by the number of myonuclei. (C) Basal (state 1 respiration) oxygen consumption rate and mitochondrial H_2_O_2_ production from hindlimb skeletal muscles 7 days following HLI using Oroboros Oxygraph-2k. (D) Representative images of Western blot analyses 7 days and 14 days following HLI for mitochondrial ETC complex I (NDUFB8-subunit), complex II (SDHB-subunit), complex III (UQCRC2-subunit), complex IV (MTCO1-subunit), and complex V (ATP5A-subunit), and 14 days following HLI for SOD2, SOD1, and myoglobin. n=3, ^*^*p*<0.05, ^**^*p*<0.01 compared to control.

To test whether this increase in mitochondria was due to a compensatory response to prior mitochondrial dysfunction, we performed functional analysis and measured levels of various proteins associated with mitochondria at days 7 and 14. We analyzed the function of isolated mitochondria at day 7 by evaluating basal (state 1) oxygen consumption rate, which did not indicate a statistical difference. However, the state 1 release of mitochondrial reactive oxygen species (ROS), H_2_O_2_, was ~3-fold higher in ischemic muscle (Fig. 4C), indicative of oxidative stress and defective mitochondria [37]. To further assess mitochondrial function, Western blot analyses (Fig. 4D) for the electron transport chain complexes also revealed that subunits of complexes I, II, and IV at day 7 and complex II at day 14 were significantly declined, suggesting potential disruption in mitochondrial bioenergetics (Supplemental Fig. S3A, B).

Western blots for antioxidant enzymes SOD2 (MnSOD), found in the mitochondrial matrix, and SOD1 (CuZnSOD), localized in the intermembrane space and cytosol [38], from day 14 homogenate were also performed as SOD is the primary defense mechanism against the superoxide ROS by converting it into the less reactive H_2_O_2_. Western blot results exhibited no change in SOD2 content, but a significant decrease in SOD1 content (Supplemental Fig. S3C). Finally, the amount of the oxygen-binding protein myoglobin, typically correlated with mitochondria content [39], was also decreased at day 14, supporting the loss of myoglobin-rich type I fibers following ischemia and suggesting decreased oxygen utilization in ischemic muscle (Supplemental Fig. S3C).

## 4. Discussion

In the present study, we report hindlimb ischemia (HLI)-induced remodeling of various niche components of the muscle stem cell (MuSC), particularly the neuromuscular junction (NMJ) of the motor unit and myonuclear and mitochondrial content of the myofiber. Due to the delayed regenerative response of skeletal muscle as a result of the abnormal perfusion to the tissues, we observed incomplete regeneration of the MuSC niche components for up to 56 days. Specifically, myofibers were still centrally nucleated while NMJs were incompletely reinnervated for at least 56 days. Ischemia-induced denervation at early time points resulted in a subsequent reorganization of the motor units that was associated with a loss of slow-twitch, type I fibers at days 14 and 28. Meanwhile, the increase in subsynaptic nuclei that maintain the NMJ synaptic transmission suggests that MuSC play a role in the repair of the NMJ. In parallel, the MuSC-mediated increase in number of myonuclei within a muscle fiber and its concomitant decrease in myonuclear domain are likely a part of the repair mechanism of skeletal muscle in order to increase the transcriptional control over the myofiber. The smaller myonuclear domain following ischemia also allows greater expression of transcripts required for mitochondrial network remodeling. Although we report an increase in mitochondrial content per nucleus, we postulate that this is a compensatory effect regulated by the accreted myonuclei to overcome the preceding mitochondrial dysfunction. While the temporal sequence of events in the various MuSC niche components following ischemia have not been thoroughly investigated altogether until this point, we report early degeneration of the NMJ and dysfunction of the mitochondria after ischemic injury, followed by MuSC-mediated remodeling of myonuclei, and finally an increase in mitochondria content to support the high energy demands of the regenerating tissues.

Because the vasculature is required in biological tissues to supply key nutrients, growth factors, chemokines, and immune cells that initiate tissue regeneration [10, 11], it is evident that functional perfusion to the tissues is a key determinant in regeneration kinetics. For instance, freeze injury to the tibialis anterior (TA) muscle results in a destruction of the vascular network and a consequent delay in regeneration of the skeletal muscle compared to chemical models of injury that demonstrate less severe vascular lesions [30]. Analogously, ischemic insults, which disrupt the perfusion to skeletal muscle, delays the initial response to injury and influences the outcome of muscle regeneration, as demonstrated by the continuous regeneration 56 days following HLI. While previous reports have shown a delayed and prolonged regenerative response to ischemia through centrally nucleated fibers up to day 14 [12], accreted myonuclei up to day 14 [40], and denervation up to day 28 [20], our data indicates that the regeneration of these niche components are sustained for at least 56 days. Nonetheless, in order to discern when these ischemic tissues are fully repaired, further testing with extended time points is required. It is noteworthy, however, that these studies, including ours, were conducted on *C57BL/6J* mice, which have been shown to exhibit more pre-existing collateral vessels [41], greater angiogenic potential, higher revascularization rates, and enhanced skeletal muscle regeneration compared to other strains of mice (i.e. *BALB/c*, *129S2/Sv*) [42]. These differences are attributed to strain-dependent genetic variation. Additionally, *BALB/c* strains of mice display a delay in MuSC activation following crush injury [43], impairing their myogenic potential. Because similar genetic polymorphisms may manifest in the population of human CLI patients, it is worthwhile to study the effects of ischemia in various mouse strains to provide a more representative animal model of CLI despite genetic differences.

In spite of the resilience to ischemic injury in *C57BL/6J* mice, we observed substantial remodeling of the myofiber and its contents, albeit with high variability in myonuclei number and mitochondrial content among different myofibers from each sample. This variability among myofibers indicates that the regenerative response occurs on a fiber-to-fiber basis likely dependent on fiber type. While we noted a loss of slow type I fibers, it is unknown whether this is a cause or effect of reorganization of the various motor unit types. It is plausible that the fiber type shifting signifies the inability of slow, oxidative motor units to reinnervate muscle fibers. Since fiber type is in part determined by motor unit type [33], slow, oxidative motor units that rely on aerobic respiration may be so severely debilitated by ischemia that they are unable to reinnervate any myofibers, resulting in the loss of type I myofibers. Alternatively, myofibers have been demonstrated to provide retrograde signaling to motor units to influence the phenotype of the nerve that innervates them [44]. Therefore, ischemia may directly induce necrosis of the oxidative type I myofibers, which thereby results in a loss of slow, oxidative motor units. To elucidate the mechanism behind the fiber type shifting and corresponding remodeling of the motor units, we plan to investigate regeneration of the slow motor neurons by staining the synaptic vesicle protein, SV2A, selectively localized in motor nerve terminals of slow, oxidative motor units [44]. Moreover, fiber type has also been implicated in determining the remodeling capacity of myonuclei. It has been found that the change in myonuclear domain is fiber-dependent, with type I fibers increasing myonuclear domain size while type IIB and IIX decrease myonuclear domain following denervation [45]. With the loss of type I fibers in ischemia, it is likely that the majority of myofibers remaining (types IIA, IIB, IIX) decrease their myonuclear domain in response to ischemic insults. Interestingly, oxidative type I fibers have also been shown to be more protected against oxidative stress damage than glycolytic type IIB/IIX fibers in acute ischemia-reperfusion involving a tourniquet [46]. However, since our findings in the more severe form of ischemic injury demonstrate a loss of type I fibers with no change in type II fibers, the type II fibers may have a more robust regenerative ability following atrophy in a chronically ischemic environment.

This altered regenerative capacity of fast type II fibers may be attributed to a change in mitochondrial dynamics. The mitochondrial network of slow type I fibers are elongated and span several sarcomeres, associated with high mitochondrial fusion and fission rates. In contrast, mitochondria from fast type II fibers are compartmentalized into columns and networks are rarely elongated [24]. In our study, we observe this elongated architecture of the mitochondrial reticulum throughout all ischemic myofibers at 7 days and between the centrally localized nuclei at 14 and 28 days following CLI, suggesting that type II fibers may adopt this feature of high fusion and fission rates as a regenerative response to ischemia. Furthermore, despite the loss of mitochondria-rich type I fibers following CLI, other researchers have demonstrated an increase in muscle mitochondrial DNA (mtDNA) in PAD patients [47], substantiating the increase in mitochondrial content at day 28. While some proteins of the mitochondria are encoded by mtDNA, the majority of mitochondrial proteins are nuclear-encoded [36], illustrated by the high mitochondrial density localized adjacent to myonuclei. There are several possible explanations for the increase in mitochondrial content. First, because of the mitochondrial dysfunction and impaired bioenergetics [48], myonuclei may respond as an adaptive measure to increase functional mitochondria to meet the high energy demand of skeletal muscle. This is accomplished by decreasing myonuclear domain size so that myonuclei can more effectively generate transcripts for biogenesis of the surrounding mitochondria. Second, oxidative stress may directly upregulate the genes required for mitochondrial proliferation [49]. Finally, the increased mitochondrial content could be a result of lack of autophagy to remove the accumulation of oxidatively damaged mitochondria [50]. To gain insights into the mechanism behind the mitochondrial biogenesis, we plan to further examine the roles of oxidative stress and autophagy as well as mitochondrial dynamics.

Additionally, our data contributes insight into ischemia-induced mitochondrial dysfunction. While the reactive oxygen species (ROS) hydrogen peroxide (H_2_O_2_) is required for redox signaling mechanisms and regulation of transcription factors, an imbalanced homeostasis resulting in excess H_2_O_2_ production can cause cellular damage by oxidation of lipids, proteins, and nucleic acids via Fenton reaction to HO• radical [51]. Mitochondria are one of the major sources of ROS and generate H_2_O_2_ by converting the reactive superoxide into the more stable H_2_O_2_, catalyzed by the antioxidant enzyme superoxide dismutase (SOD). To elucidate the precise origin of increased H_2_O_2_ production in ischemic mitochondria, enzymatic activity of the mitochondrial membrane complexes and isoforms of SOD as well as quantification of superoxide levels is required. While complex I releases ROS into the mitochondrial matrix, complex III releases ROS to both sides of the inner membrane [52]. Since the content of SOD2 (MnSOD), primarily localized in the matrix, is unchanged while the content of SOD1 (CuZnSOD), found in the intermembrane space and cytosol, is decreased, we hypothesize that ROS are present in the intermembrane space due to complex III-mediated electron leak. Though we acknowledge that our analysis of mitochondrial function is not fully comprehensive, we plan to analyze complex III activity in order to test this hypothesis.

Finally, although increased H_2_O_2_ alone does not impact muscle stem cell number, it has been shown to limit their myogenic potential [53]. However, we present clear evidence of increased MuSC frequency and MuSC-mediated expansion of myonuclei 7 days following ischemia, despite the greater H_2_O_2_ emission from mitochondria. While much of the analyses on muscle stem cells in ischemic environments performed *in vitro* have been controversial [54, 55], they lack the niche components of the MuSC and only provide a myopic understanding of muscle stem cell behavior. Additionally, *in vivo* studies that report a decrease in MuSCs following ischemia have used Myf5 as a marker for muscle stem cells [12, 56], expressed in activated MuSCs. These experiments could be improved upon as they do not analyze the self-renewal of the quiescent Pax7^+^ MuSC population or differentiation and fusion of the MuSC into the myofiber. Furthermore, because the number of myonuclei, which are derived from MuSCs, are a causative factor in determining myofiber size [57], the accumulation of myonuclei likely assists in the regeneration of atrophied myofibers. Accretion of myonuclei allows for subsequent hypertrophy following the ischemia-induced atrophy so that the myonuclear domain and myofiber volume can return to normal once perfusion is improved at day 28. The accreted myonuclei may also explain the increased subsynaptic nuclei that assist in the regeneration of the NMJ. As MuSCs play a vital role in NMJ regeneration [22], we suspect that this expansion of subsynaptic nuclei are also mediated by proliferation and differentiation of the muscle stem cell. To further validate these results, we plan to conduct a loss-of-function study by analyzing myonuclei number and subsynaptic nuclei number in transgenic mice with depleted MuSCs. However, it is important to note that age is an etiologic factor for CLI and the regenerative potential of muscle and MuSC frequency both decline with age [58]. Therefore, the MuSC-mediated regenerative response to ischemia may be impaired in elderly patients. To address this, further examination of CLI in aged mice is required.

In summary, we demonstrate remodeling of MuSC niche components following ischemic injury, notably through neuromuscular junction repair, increased myonuclei number, decreased myonuclear domain, and greater mitochondria content per nucleus of skeletal muscle fibers. Taken together, these data indicate that CLI resets the myonuclear and mitochondrial domains as part of the regenerative mechanism in ischemic conditions. The findings from this study illustrate the complex and prolonged regenerative response of the MuSC niche following critical limb ischemia and serve as a fundamental basis to develop therapies that target the skeletal muscle in order to prevent extensive muscle damage while collateral vessels are formed.

## Acknowledgements

Research reported in this publication was supported by the National Institute of Arthritis and Musculoskeletal and Skin Diseases of the National Institutes of Health under Award Number R21AR072287 (YCJ) and grants from Regenerative Engineering and Medicine. The content is solely the responsibility of the authors and does not necessarily represent the official views of the National Institutes of Health. We also thank the Physiological Research Laboratory and core facilities at the Parker H. Petit Institute of Bioengineering and Bioscience at the Georgia Institute of Technology for the use of shared equipment, services, and expertise.

## Supplemental Information

**Supplementary Table S1.**
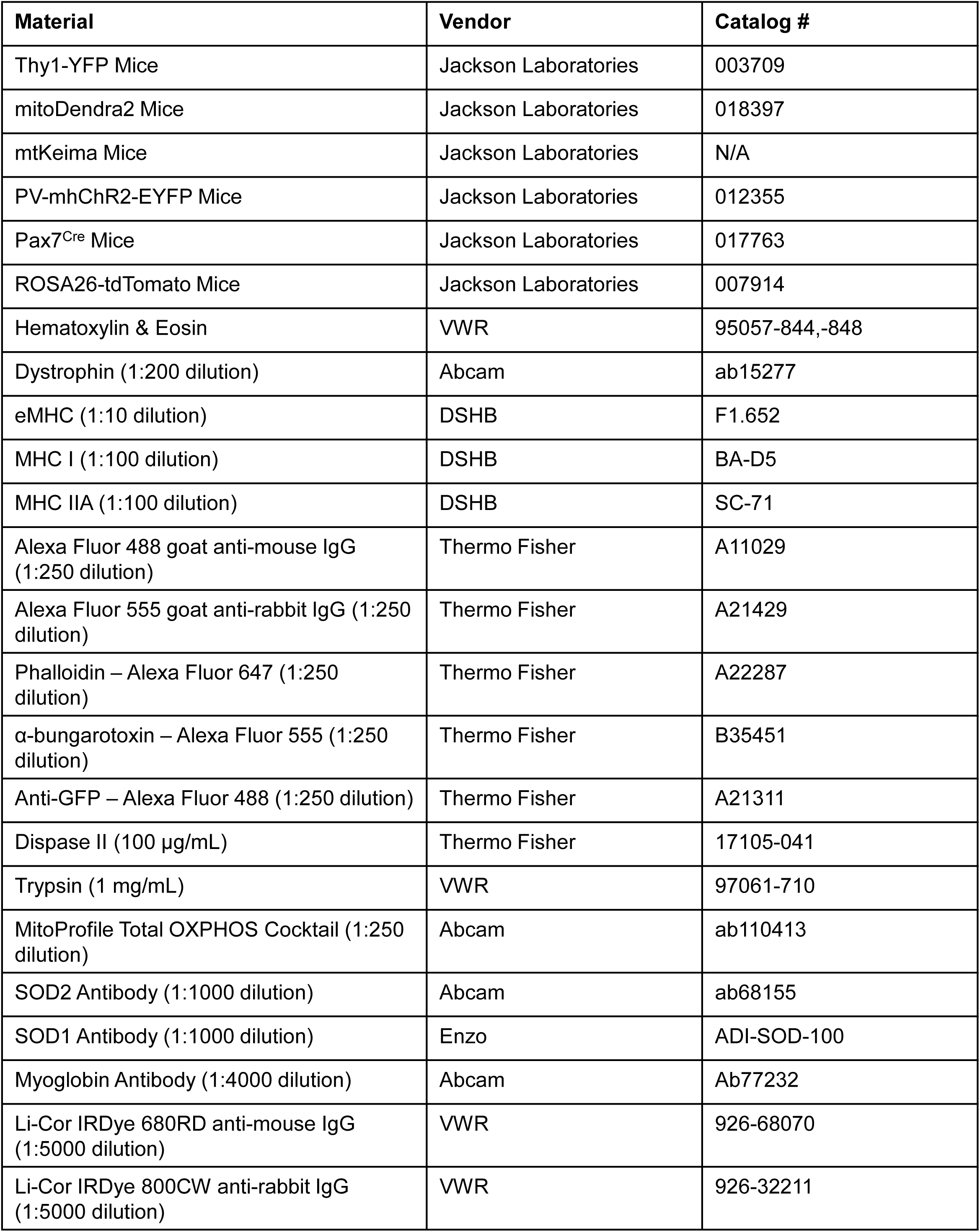
List of materials, dilution factors, vendors, and catalog numbers used in this study.

**Supplementary Table S2.**
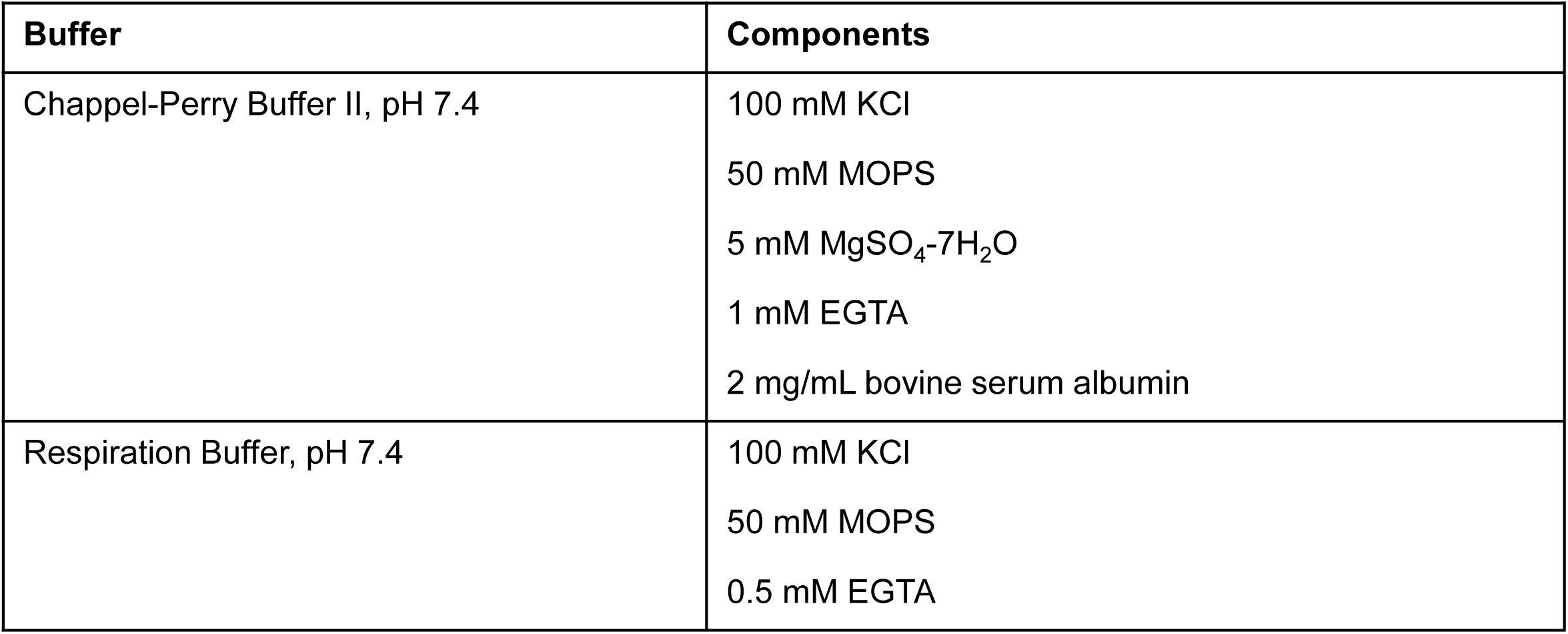
List of buffers used for mitochondrial isolation and their respective components.

**Supplementary Figure S1.**
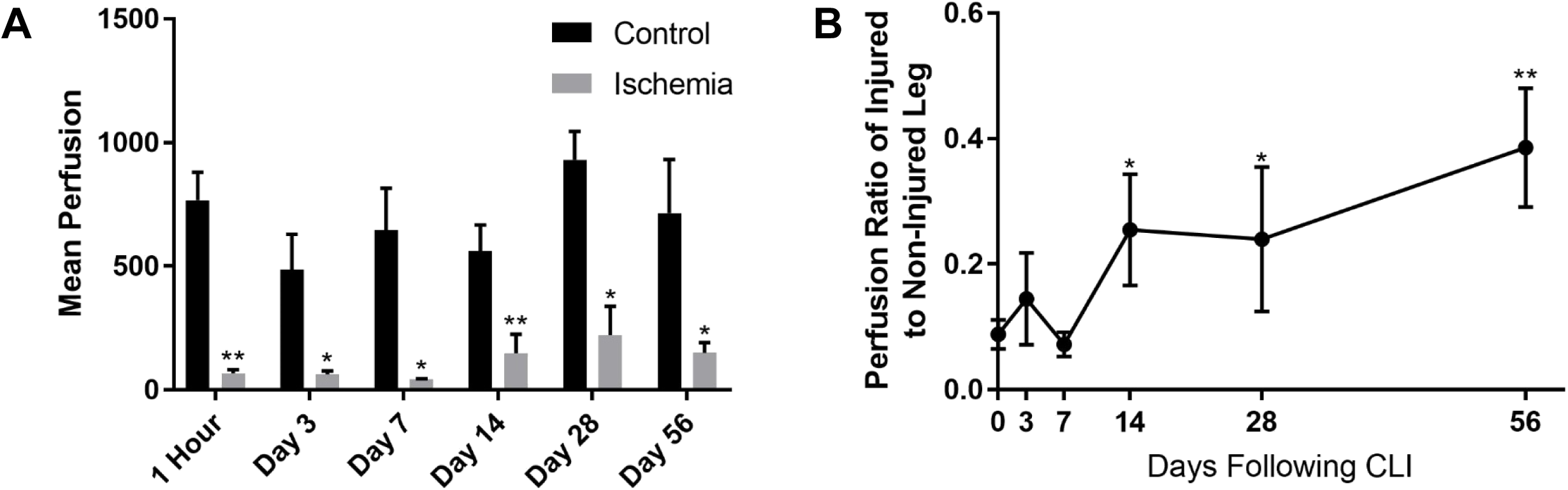
Quantification of Laser Doppler Perfusion Imaging. (A) Mean perfusion to hindlimb distal to the knee in ischemic leg and contralateral control over 56 days. n=3, ^*^*p*<0.05, ^**^*p*<0.01 using two-way ANOVA with Tukey’s*post hoc* test. (B) Ratio of perfusion in ischemic leg to control over 56 days. n=3, ^*^*p*<0.05, ^**^*p*<0.01 compared to day 0.

**Supplementary Figure S2.**
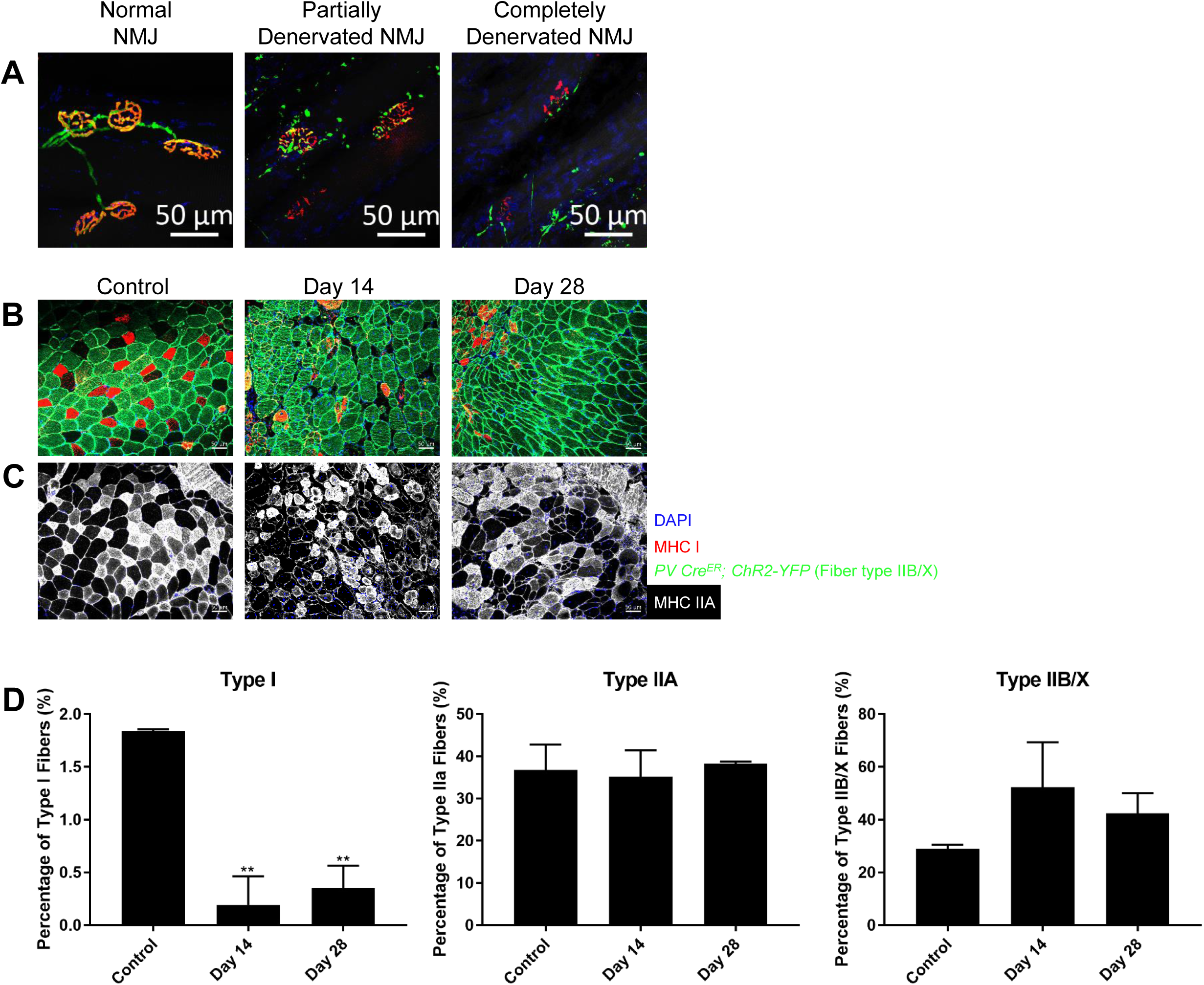
NMJ Categorization and Fiber Type Shifting. (A) Representative images of normal, partially denervated, and completely denervated NMJ. Bungarotoxin pseudo-colored in red, Thy1 in green. (B) TA cross-sections of *PV*-*Cre; ChR2*-*YFP* mice for expression of fiber type IIB/IIX and stained for fiber type I (MHC I) in control, 14 days, and 28 days following HLI. Nuclei pseudo-colored in blue, MHC I in red, MHC IIB/IIX in green. (C) TA cross-sections of wildtype mice stained for fiber type IIA (MHC IIA) in control, 14 days, and 28 days following HLI. Nuclei pseudo-colored in blue, MHC IIA in purple. Scale bars represent 50 μm. (D) Percentages of type I, IIA, and IIB/X fibers, respectively. n=3, ^**^*p*<0.01 compared to control.

**Supplementary Figure S3.**
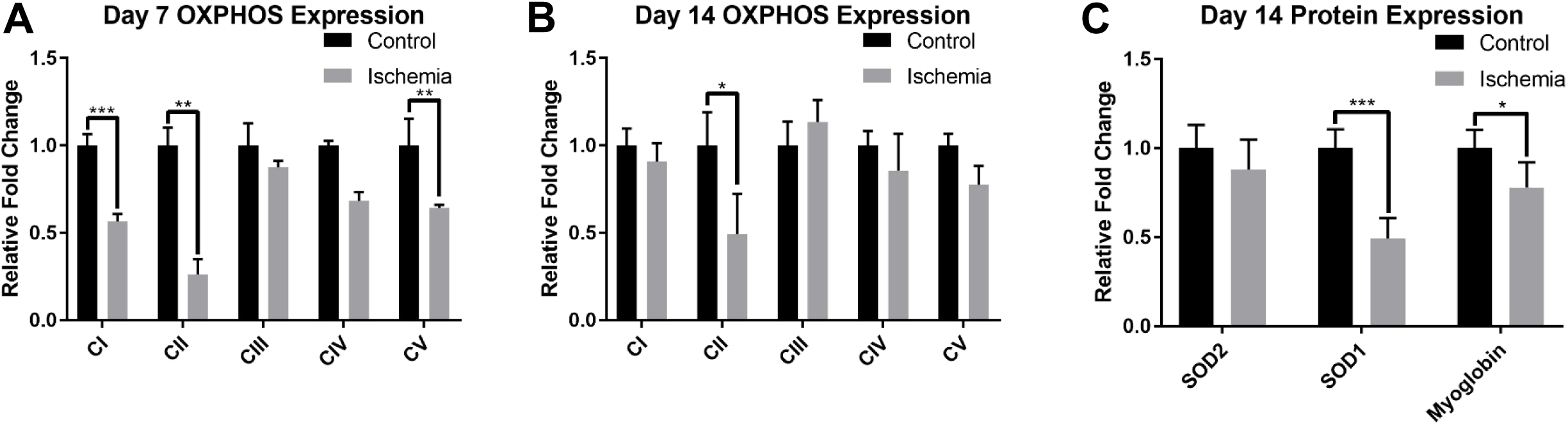
Quantified Western blot bands. (A) Relative fold change for ETC complexes 7 days following HLI. (F) Relative fold change for ETC complexes 14 days following HLI. (G) Relative fold change for SOD2, SOD1, and myoglobin. n=3, ^*^*p*<0.05, ^**^*p*<0.01, ^***^*p*<0.001 compared to contralateral control.

